# Glioblastoma cells use an integrin- and CD44-mediated motor-clutch mode of migration in brain tissue

**DOI:** 10.1101/2023.10.23.563458

**Authors:** Sarah M Anderson, Marcus Kelly, David J. Odde

**Affiliations:** Department of Biomedical Engineering, University of Minnesota, Minneapolis, Minnesota, USA

## Abstract

Glioblastoma (GBM) is an aggressive malignant brain tumor with 2-year survival rates of 6.7% [1], [2]. One key characteristic of the disease is the ability of glioblastoma cells to migrate rapidly and spread throughout healthy brain tissue[3], [4]. To develop treatments that effectively target cell migration, it is important to understand the fundamental mechanism driving cell migration in brain tissue. Here we utilized confocal imaging to measure traction dynamics and migration speeds of glioblastoma cells in mouse organotypic brain slices to identify the mode of cell migration. Through imaging cell-vasculature interactions and utilizing drugs, antibodies, and genetic modifications to target motors and clutches, we find that glioblastoma cell migration is most consistent with a motor-clutch mechanism to migrate through brain tissue *ex vivo*, and that both integrins and CD44, as well as myosin motors, play an important role in constituting the adhesive clutch.

## Introduction

Glioblastoma multiforme (GBM) is an aggressive malignant brain tumor with extremely low 5-year survival rates[1]. The current standard of care includes surgery, radiation, and temozolomide chemotherapy, though these treatments have failed to prevent recurrence and fail to result in long-term survival. One key characteristic that drives the aggressiveness of the disease is the ability of the tumor cells to migrate rapidly and spread throughout the brain[3]–[5]. Developing an in depth understanding of the mechanisms that cancer cells use to migrate could provide opportunities for developing novel strategies for limiting cancer spread.

Several models of cell migration have been proposed. The motor-clutch mechanism of force transmission describes the process of transmitting intracellular forces to the extracellular matrix (ECM). In this model, forces in the cell are generated through non-muscle myosin II motors and F-actin polymerization, leading to retrograde movement of the F-actin filaments away from the leading edge toward the cell center [6]. Adhesion molecules, or molecular “clutches”, connect the F-actin network to the ECM to transmit force to the environment[7]–[9]. The motor-clutch model simulates this behavior by describing the stochastic breakage and rebinding of clutches in response to a moving actin filament network/bundle[10], [11]. Though other versions of the model have been developed[12], the simplest version regards the F-actin network/bundle as a rigid rod, which can bind to an array of linear springs, or clutches. The clutches exhibit force-dependent unbinding and transmit force to a linear substrate spring that describes the elastic properties of the extracellular environment. Eventually, the clutches at one protrusion fail stochastically, while the clutches at other protrusions persist, thus breaking the symmetry, so that the failed protrusion becomes the trailing edge, and the persisting protrusions constitute the leading edge. This results in net movement of the cell towards the leading edge. The movement of the cell thus moves opposite of the direction of substrate deformation, which is one of the key predictions of the motor-clutch model. The trailing edge experiences pulling forces until the protrusions fail and the substrate mechanically relaxes. The behavior of the entire cell was simulated by Bangasser et al. (2017) [13], who developed a cell migration simulator (CMS), which links multiple motor clutch modules via a spring at the cell body, to stochastically simulate whole-cell migration. One key finding of this model is the biphasic relationship between cell migration speed and motor-to-clutch ratio in which balanced motors and clutches leads to optimal cell migration[13], [14].

The physical paths that glioma cells use to invade healthy tissue has been studied extensively. Post-mortem histological analysis was used to determine that glioma cells migrate along the brain parenchyma, blood vessels, white matter tracts, and the subarachnoid space below the meningeal covering of the brain[15], [16]. Several studies have used dynamic analysis in brain tissue to identify the vasculature as the primary tract of cell migration[17]–[21]. Liu et al. imaged the deformation of vasculature during migration of U251 glioma cells and found that cells primarily exert a pulling force at their leading edge, consistent with a motor-clutch model and inconsistent with other proposed modes of migration, though they analyzed only a small number of cells and one high-passage cell line[22].

Within the motor clutch framework, integrins are generally thought of as the primary cell adhesion molecule that acts as the clutch, linking the actin cytoskeleton to the extracellular matrix via the adaptor proteins talin, vinculin, and alpha-actinin [23]– [25]. Integrins are a family of adhesion proteins that bind directly to ECM proteins and play a critical role in cell migration and signaling in many cell types and in cancer[26], [27], [28]. They have a heterodimeric structure composed of alpha and beta subunits. These subunits bind directly to specific peptide sequence, such as RGD, on ECM substrates, including collagen, laminin, and fibronectin[29]. Talin, which has two human isoforms, talin1 and talin2, acts as a force-bearing connection, or adaptor, between integrin cytoplasmic tails and actin filaments. Recent studies have identified other molecules that could have the potential to act as a molecular clutch, such as the adhesion molecule CD44. CD44 binds to hyaluronic acid via its extracellular domain and connects to the actin cytoskeleton intracellularly via adaptor proteins ezrin, radixin, and moesin, the so-called ERM proteins, and ankyrin[30]–[33]. CD44 has also been implicated in mediating glioma cell migration[34]–[36]. Klank et al. found a biphasic relationship between survival in both humans and mice and CD44 expression, where an intermediate expression of CD44 correlated with fastest migration in mice and lowest survival in both mice and humans, as predicted by the CMS[37]. This study also demonstrated that natural variations in expression of CD44 across subtypes biphasically correlates with patient survival. Specifically, the proneural subtype was shown to have lower CD44 expression and the mesenchymal subtype was shown to have higher CD44 expression, with the former subtype exhibiting modestly longer survival. These findings have been supported by other studies demonstrating that glioblastoma cells can migrate through hyaluronic acid-rich matrices via CD44 adhesion [36] and that cells from mesenchymal glioma tumors have higher CD44 expression and migrate faster than their proneural counterparts [38].

However, the universality of the motor-clutch model has been questioned. In particular, Lämmerman et al. showed that dendritic cells in 3D could migrate without integrins[39], a key component of the molecular clutch. However, there are potential limitations of this finding, in particular not all integrins were knocked out, and other adhesion molecules that could mediate motor-clutch migration, such as CD44, were presumably still active. It is worth noting that a high copy number of relatively weak, i.e. non-specific, adhesive bonds could still serve the function of a clutch in the motor-clutch framework. However, as a result of this finding, alternate models of cell migration were proposed. Stroka et al. proposed the osmotic engine model, in which cell migration is driven by a spatial gradient of ion/osmolyte pumps in the plasma membrane that lead to directed water fluxes throughout the cell [40]. In this model, migration does not require on adhesion to the environment. Instead, the cell establishes a spatial gradient of ion pumps that creates a net influx of ions at the leading edge and net outflux of ions at the trailing edge. This creates an electro-osmotic gradient across the cell cytoplasm, driving water to permeate into the cell at the leading edge and out of the cell at the trailing edge, leading to net movement of the cell forward. Following this hypothesis, Stroka et al. found cell migration is sensitive to changes in Na+/H+ exchangers, changes in aquaporin water channels, specifically AQP5, and osmotic shock. However, their experiments were conducted in confined microchannels in vitro, and whether this mode of migration occurs in physiological relevant environments, such as for glioblastoma cells in brain tissue, is an open question [40]. A cell migrating under this mechanism could still maintain adhesion to the environment, so as the osmotic forces propel the cell forward adhesions would presumably exert a frictional force resisting the motion, and so blocking adhesions would result in an increased cell migration speed. In addition, the cell would push the environment away from the leading edge, which is inconsistent with the small set of observations from Liu et al. for U251 cells in normal mouse brain slices. However, again, these results were based on a limited number of observations for a highly passaged cell line, and it is unknown whether lower passage glioblastoma cells use the osmotic engine model for migration in brain tissue.

Another mechanism of cell migration that has been proposed is bleb-based motility[41]. A variety of cell types have been shown to produce blebs *in vitro*, including A375 melanoma cells, breast cancer cells, and T cells[42]–[45]. The most *in vivo*-like environment blebbing has been observed in is 3D collagen gels[44], [46]. A bleb is a local transient loss of plasma membrane to F-actin cortex cohesion that causes a plasma membrane-bounded blister-like protrusion to rapidly extend from the cell[46], [47]. Blebbing enables the plasma membrane to rapidly advance, faster than F-actin can polymerize. Subsequently, the actin-myosin cortex reforms under the plasma membrane, adhesions can form, and the intracellular forces can be transmitted to the environment. In order for the cell to migrate following the rapid extension of the plasma membrane, adhesions must presumably form at the leading edge and engage the cell’s motor-clutch modules, pulling the cell forward before the bleb can be retracted. In this description, bleb-based motility can be regarded as a form of motor-clutch-mediated migration, where the plasma membrane advances by blebbing rather than F-actin self-assembly, as in the original motor-clutch model. Even so, during bleb extension, this model predicts that the extracellular environment will be pushed away from the leading edge, whereas the F-actin assembly-based motor-clutch model predicts only pulling forces at the leading edge.

In the present study, the migration of a high passage gliomblastoma cell line, U251, and 6 low passage patient-derived xenograft (PDX) glioblastoma lines were investigated in mouse brain-slice organotypic culture to discern the cell migration mechanism in brain tissue *ex vivo*. Brain slices have been found to preserve the cytoarchitecture of the brain and additionally be an effective way to study the effects of small molecule inhibitors on cell migration [48], [49]. Traction dynamics and cell migration speeds in mouse brain slices were assessed using two-color swept-field confocal imaging with glioma cells either expressing GFP-actin or a green membrane dye and the brain vasculature labeled with a red fluorescent marker (isolectin-b4-rhodamine). Genetic modification and drug perturbations were used to target specific motor and clutch molecules potentially involved in migration. We find that the observed traction dynamics are consistent with glioma cells using a motor clutch mode of migration and argue against other models, including the osmotic engine model and bleb-based motility model. We also find myosin, integrins and CD44 to be important for cell migration, with implications for therapeutic targeting of glioblastoma cell migration.

## Materials and Methods

### U251 cell culture

The U251 human glioblastoma cell line was obtained from Dr. G. Yancey Gillespie (University of Alabama at Birmingham) and was authenticated using the short tandem repeat assay (ATCC) in July 2023. Stably transfected U251-GFP-actin cells were used[13]. Cells were cultured in T25 plastic tissue culture flasks (353108; Becton Dickinson, Franklin Lakes, NJ) in a humidified 37°C, 5% CO_2_ incubator. Dulbecco’s modified Eagle’s medium/F-12 (31765-035; Gibco, Gaithersburg, MD) supplemented with 8% fetal bovine serum (FBS; 10438-026; Gibco) and 1x penicillin/streptomycin (100 IU/mL Penicillin, 100 μg/mL Streptomycin; 30-001-CI; Corning, Corning, NY) (P/S) was used to culture the cells. Before plating cells onto brain slices for migration studies, cells were removed from the flask using a 5 minute incubation with 0.25% trypsin with EDTA in Hanks’ balanced salt solution (25-052-CI; Gibco).

### Mayo PDX cell culture

PDX lines were obtained from the Mayo Clinic Brain Tumor Patient-Derived Xenograft National Resource[50]. Cell lines that were frozen in FBS-containing media (GBM44, GBM85, GBM80) were cultured in T25 plastic tissue culture flasks (353108; Becton Dickinson, Franklin Lakes, NJ) in a humidified 37°C, 5% CO_2_ incubator with Dulbecco’s modified Eagle’s medium/F-12 (31765-035; Gibco, Gaithersburg, MD) with 8% fetal bovine serum (10438-026; Gibco) and 1x penicillin/streptomycin (30-001-CI; Corning, Corning, NY). Cells that were frozen without FBS (GBM12, GBM117, GBM39, GBM85) were cultured on T25 plastic culture dishes coated in 10% Matrigel with DMEM/F12 with 1x B-27 supplement (12587010; Gibco) and 1x P/S.

### Creation and maintenance of cell lines

CD44 knockout (KO) was achieved as described previously [51]. TLN1 KO was achieved using the CRISPR/Cas9 system. A guide RNA (sequence AACUGUGAAGACGAUCAUGG) was created to target TLN1. U251 cells were transfected with Cas9 nuclease and gRNA using the Neon Electroporation system (Voltage = 1150V, Pulse width = 30ms, number of pulses = 2, interval between pulses = 1 ms). Serial dilution was then used to generate several hundred single cell clones. Amplification and sequencing of the TLN1 gene for some of the single clones was used to identify 4 clones with a homozygous TLN1 KO. Western blot was used to confirm knockout of TLN1 and verify that TLN2 expression was not upregulated in response to TLN1 KO (Supplementary Figure 2).

### Preparation of mouse brain slices

All animal treatments and experiments were conducted in accordance with Institutional Animal Care and Use Committee at the University of Minnesota approved protocols. Adult B57BL/6 mice aged 8-12 weeks from the Jackson Labs were terminally anesthetized in a CO2 chamber and then perfused transcardially with ∼20mL isotonic saline until the liver appeared tan colored. The brains were extracted and transferred into chilled oxygenated artificial cerebrospinal fluid (124 mM NaCl, 2.5 mM KCl, 2.0 mM MgSO_4_, 1.25 mM KH_2_PO_4_, 26 mM NaHCO_3_, 10 mM glucose). The cerebrum was sectioned into 300 μm coronal slices using a vibratome at 80 Hz vibration speed, 2.5mm amplitude, and 0.5mm/sec advance speed (Lafayette Instruments; 7000SMZ-2).

### Organotypic brain-slice coculture with glioma cells

Brain slices were transferred into DMEM/F12 media containing 8% FBS and 1x P/S and stored in an incubator at 37°C and 5% CO_2_. Two hundred thousand cells were plated onto a slice and incubated overnight to allow cells to infiltrate into the slice. Cells typically invade up to 100 μm into the tissue. Isolectin GA-IB4 (Alexa Fluor 568; Molecular Probes, Eugene, OR) was added to the slice to stain the vasculature at least 30 minutes before imaging. The slice was transferred into a 6-well glass-bottom 35 mm culture dish (P35G-0-20-C; MatTek, Ashland, MA) with fresh media. A tissue culture anchor (SHD 42-15; Warner Instruments, Hamden, CT) was placed on top of the brain slice to prevent it from drifting during imaging.

### Brain slice incubation with drugs and antibodies

For experiments that included drug targeting, the drug was added to the dish containing the brain slice and cell co-culture 3 hours before imaging to allow the drug to diffuse into the slice. When the slice was moved into fresh media for imaging, drug was added to keep the concentration consistent. For experiments with α-CD44 MAB, the slice and cells were incubated with α-CD44 mAb for 30 minutes before cell plating.

### Confocal imaging of brain slices

For experiments with brain slices, the slice was imaged using a Zeiss LSM 7 Live dual-color swept-field laser confocal microscope (Carl Zeiss, Oberkochen, Germany) with a 20x 0.45 NA objective lens capable of simultaneous imaging in both the green (GFP) and red (IB4) channels as previously described[22]. Maximal intensity projections from multiple z-stacks were used to generate 2D images for quantitative shape and motion analysis. A 3×3 tile scan was imaged, and the number of z-stacks was adjusted to ensure that the data acquisition of the whole slice was completed in under 10 min (eight to nine planes with 10 μm separation were typically used). The z-stacks were then imaged every 10 min for up to 10 h at 37°C in a humidified 5% CO_2_ environment.

### Brain slice registration and cell tracking

To track single cell migration, relevant z planes were first selected to exclude cells migrating outside of the brain slice, particularly on the glass. Maximum intensity projections from these z-stacks were used to assess single cell migration speeds. Stage drift and tissue relaxation were registered using an affine transformation in MATLAB’s imregister_tform function. Single cells were then tracked using ImageJ’s TrackMate with radius of 10-15 μm and threshold of 2-10 μm. The random motility coefficient (D) was calculated by linear fitting to the mean-squared displacement (MSD) versus time (t).

### Motor clutch modeling

The motor clutch model as described by Chan & Odde [10] and Bangasser et al.[11] was used to simulate substrate load-and-fail dynamics and output plots of substrate displacement versus time. A custom Matlab code was developed to isolate peaks in displacement and fit a line to the loading of the substrate, which estimates the stretching rate of the substrate.

### Stochastic cell migration simulator

The cell migration simulator as described by Bangasser et al.[13] and Klank et al.[37] was used to simulate 10 migrating cells under various numbers of motors and clutches for 6 hours each. The simulation parameters are shown in Table 1.

**Table 1.** Motor clutch model (MCM) with Kelvin-Voigt substrate parameters.

|  |  |  |  |
| --- | --- | --- | --- |
| $n_m$ | Total number of motors | 50 | |
| $n_c$ | Total number of clutches | 50 | |
| $k_s$ | Substrate spring constant | variable | pN/nm |
| $c_s$ | Substrate damping coefficient | variable | pN/nm |
| $F_m$ | Myosin motor stall force | 2 | pN |
| $v_u$ | Unloaded actin flow rate | 120 | nm/sec |
| $k_{on}$ | Clutch on-rate constant | 1 | $\text{sec}^{-1}$ |
| $k_{off}^*$ | Unloaded clutch off-rate | variable | $\text{sec}^{-1}$ |
| $k_c$ | Clutch spring constant | 1 | pN/nm |
| $F_b$ | Characteristic clutch rupture force | 2 | pN |

### 2D polyacrylamide gel synthesis

Polyacrylamide hydrogels (PAGs) were cast onto No. 0 glass bottom 35 mm culture dishes (MatTek P35G-0-20-C) using a previously described method and formulation [13], [52]. Cast gels were then coated using sulfo-SANPAH (Thermo Fischer Scientific, Waltham, MA) as previously described (Bangasser et al., 2017; Wang and Pelham, 1998). In this study, Type I Collagen (354236, Corning, Corning, NY) or anti-CD44 antibody α-CD44 MAB (BDB553131, BD Biosciences, San Jose, CA) were used to coat PAGs. In general, 200 μg/mL Col I solution was used and 1-300 μg/mL α-CD44 MAB solution was used depending on the experiment. In addition, 0.7, 4.6, 9.3, 19.8, 98.5, 195 kPa stiffnesses were used in this study, which were obtained by varying the cross-linker and polymer concentrations as previously described[13].

### Statistical analysis

Statistics were completed in PRISM using a Kruskal-Wallis/Mann-Whitney test with multiple comparisons to account for the number of datasets that were being compared.

## Results

### Human glioblastoma cells exert pulling forces on brain vasculature

Glioma cells, including high passage U251 cells and patient derived xenograft (PDX) lines from the Mayo Clinic, were plated on 300 μm thick normal mouse brain slices. The cells, which were stained to be green fluorescent, adhered to and migrated into the brain slice and were imaged inside the slice using confocal microscopy. This method allowed us to both assess single-cell migration speed and investigate interactions between cells and the vasculature. As described by Liu et al., the direction and speed that cells are deforming vasculature can be determined by generating a kymograph of the leading edge of the cell. Figure 1A shows example kymographs for two scenarios: motor-clutch-predicted pulling and osmotic engine-predicted pushing. Fig. 1B-E shows examples of single cells from PDX lines and U251 cells migrating along vasculature. Cells can be seen pulling the blood vessel towards itself while at the same time moving in that direction. This behavior can be captured by generating a kymograph of the leading edge. Figure 1 F-I shows kymographs for 4 cells pulling on the vasculature. Green lines show the movement of the cell and magenta lines show the movement of the vasculature. These examples show the movement of the blood vessel towards the cell, opposite the direction the cell is moving, as is consistent with a motor-clutch mode of migration but inconsistent with an osmotic engine mode of migration. The angle of the vascular movement in the kymograph, marked by a red dotted line, can be used to quantify the speed at which the vasculature is being deformed. This process was repeated for many cells, including both U251 and a variety of PDX lines. While the majority of cells are either not adhered to vasculature or show no deformation, of the cells visibly interacting with vasculature, 14 cells from 4 different PDX lines (GBM39, GBM44, GBM85, GBM80) and 8 U251 cells were analyzed and all deformations but one were consistent with a motor-clutch mode of migration (Fig 1J). The mean maximum speed of deformation was 2.2 with s.e.m. 0.52 nm/sec, well within the capabilities of myosin motor sliding velocities, which have been found to be around 3 μm/sec [53]. Observed rates of deformation would be expected to be significantly smaller than maximum myosin motor sliding velocities because the actin filaments experience resistance from the bound adhesion molecules, which we next modeled using a motor-clutch model. Overall, these results are consistent with a motor-clutch model for cell migration, and inconsistent with an osmotic engine model for cell migration.

**Figure 1.**
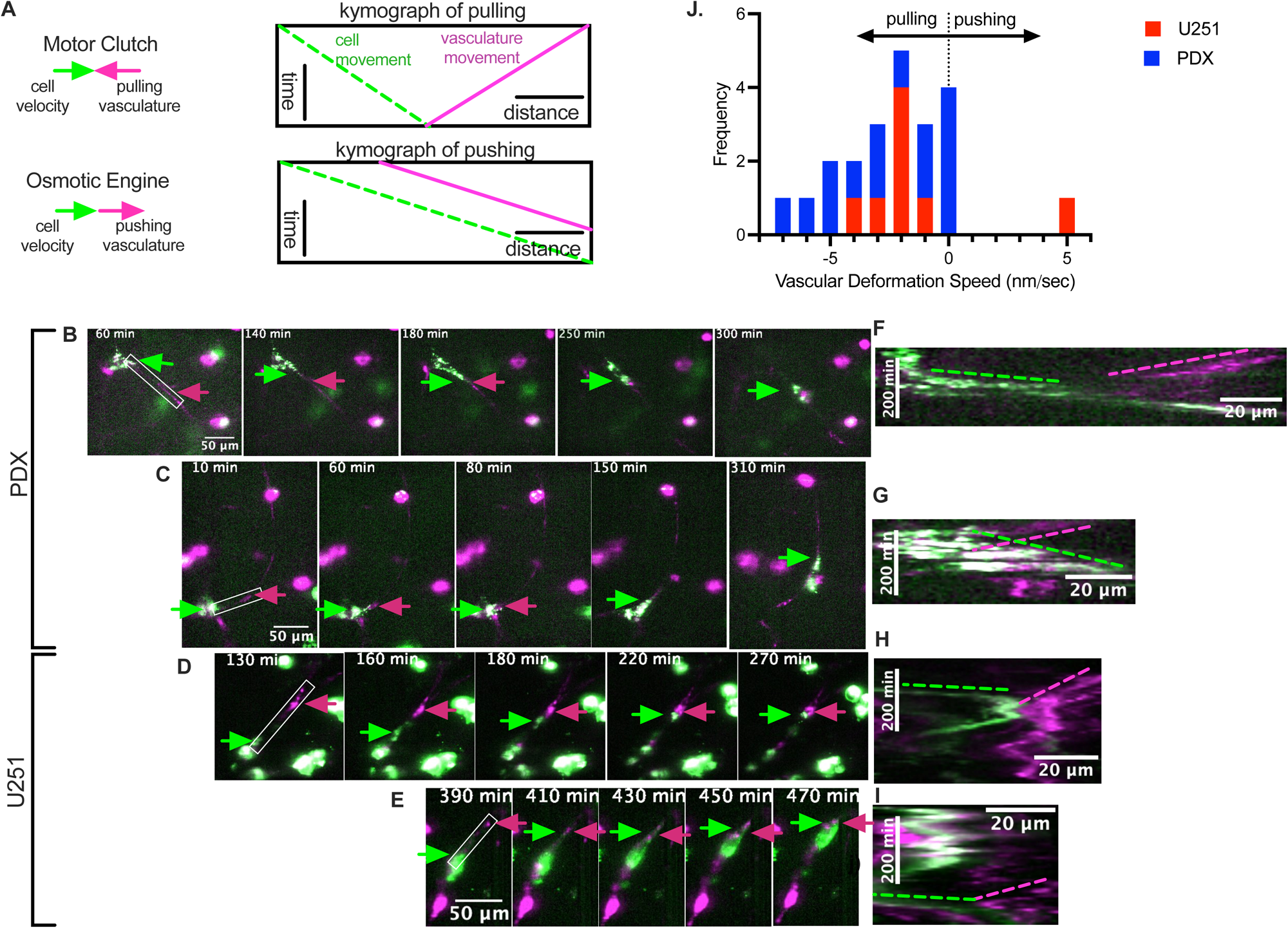
Glioblastoma cells exert pulling forces during migration that are consistent with a motor-clutch model for migration. A) Schematic depicting example kymographs of a cell pulling on vasculature vs. pushing on vasculature. Examples of B-C) Mayo PDX cells (GBM44) and D-E) U251 cells (green) imaged by confocal microscopy migrating along blood vasculature (magenta). Green arrows indicate the leading edge of the cell and magenta arrows indicate the vasculature being deformed. The cell can be seen pulling the vasculature towards itself as it moves forward. F-I) Kymographs of the leading edge of the cell as denoted by the white box on a-d showing motion of the vasculature towards the cell. j) Calculated rates of vasculature movement show that for PDX and U251 cells the vascular is pulled towards the cell at 0-7 nm/sec with one U251 cell observed to push the vasculature away. The mean speed of vasculature deformation is 2.2 nm/sec with s.e.m. 0.52 nm/sec (n=22 observations).

### Motor clutch model predicts cell traction dynamics on vasculature

Under a motor clutch mode of migration, F-actin retrograde flow applies inward forces to the environment, i.e. towards the advancing cell. When the cell moves forward and the trailing edge releases, there would be a pulling at the leading edge and relaxation at the trailing edge. By contrast, in an osmotic engine model, the cell would be moving forward independent of its adhesions, applying frictional forces to the surrounding environment. This would result in a *pushing* at the leading edge and a pulling at the trailing edge. Thus, the directions of the forces at the leading edge in the two models are in opposite directions, providing an excellent opportunity for model discrimination, as summarized above and in Fig. 1.

The motor clutch model with the substrate modeled as a Kelvin-Voigt (KV) material [54] (Fig 2A) was used to simulate deformation of the substrate in response to cellular forces. A KV model was used because the simplification of the substrate to a simple Hookean spring could not achieve substrate displacements as large as what was observed experimentally due to load and foal of the substrate spring, while a KV model allows the substrate to deform further. Additionally, with the substrate modeled as a simple Hookean spring, the displacement must snap back to its original state, which was not always observed experimentally. A KV model can describe plateauing of the displacement. Adebowale et al., previously used a Standard Linear Solid (SLS) model for viscoelasticity, but the KV model is a special case of the SLS model where one spring is considered infinitely stiff, and is therefore simpler[55]. Model parameters are shown in Table 1. The substrate deformation can be plotted over time in response to changing various motor-clutch parameters. The stiffness of the substrate spring determines the maximum displacement to which the substrate can be stretched (Fig 2B). A magenta dashed line indicates the average stretching speed of 2.2 nm/sec that was found experimentally. The damping constant impacts the rate at which it approaches the maximum displacement (Fig 2C). Altering a parameter that impacts the strength of the clutches, including number of clutches, characteristic clutch bond force, and unloaded off-rate of the clutches, will also change the rate at which the substrate approaches maximum deformation. In Figure 2D, the clutch unloaded off-rate constant is varied (k_off_). In order to simulate deformations of 20-40 µm at a speed on the order of 1 nm/sec, small spring constants (0.01-0.001 pN/nm) and relatively large damping constants (10-60 pN/nm) were used. Pogoda et al. analyzed the time-dependent viscoelastic properties of brain[56]. Though they did not analyze storage and loss moduli at frequencies as low as 10^-3^-10^-4^ Hz, the trends in their data would imply that at these frequencies, the viscous loss modulus would be significantly higher than the elastic storage modulus, as was found to be the case here. Thus, inward deformations of the vasculature at the leading edge observed experimentally can be computationally reproduced in a motor-clutch model assuming a Kelvin-Voigt model for local brain-vasculature mechanics. These experimental and computational results together support the hypothesis that cells migrate using a motor clutch mode of migration and contradict the hypothesis that cells migrate using an osmotic engine mode of migration.

**Figure 2.**
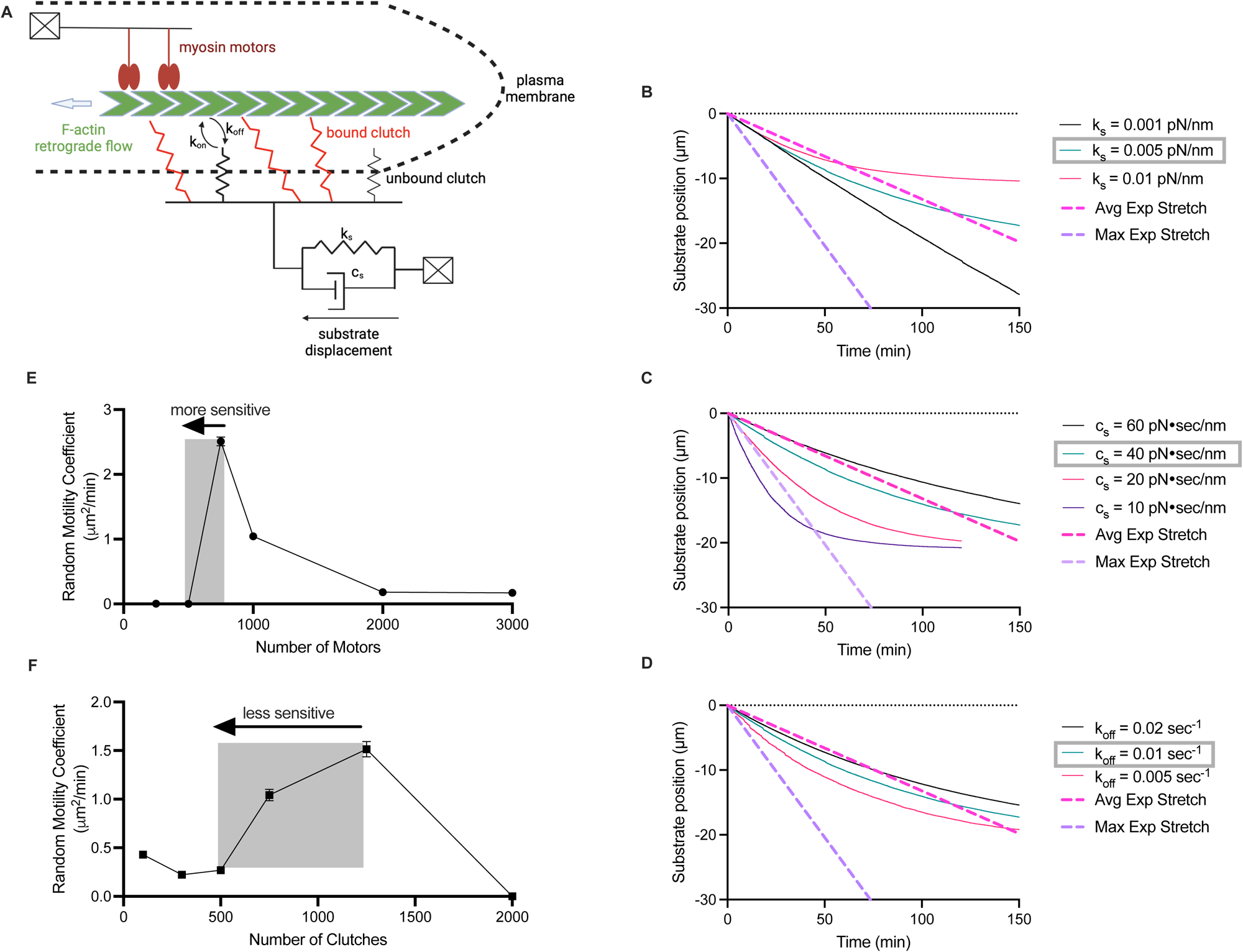
Motor clutch model and cell migration simulator predict cell traction and migration behavior in brain slices. A) Schematic of the motor clutch model with the substrate modeled as a Kelvin-Voigt viscoelastic material. Displacement of the substrate simulated over time in response to changes in B) k_s_, the substrate spring constant, C) c_s_, the substrate damping constant, and D) k_off_, the unloaded clutch unbinding rate. A dashed magenta line indicates the mean speed of vascular deformation found experimentally and a dashed purple line indicates the maximum speed of vascular deformation observed experimentally. The cell migration simulator was used to model the relationship between E) number of motors and F) number of clutches and RMC. Cells are more sensitive to decreases in motor number below the optimum than clutch number below the optimum.

### Cell Migration Simulator predicts response to motor and clutch targeting drugs

To further test whether cells migrate via a motor clutch mode of migration in brain slices, we first sought to understand theoretically how migration would be affected by drugs targeting motors and clutches is important. The cell migration simulator, described by Bangasser et al., was used to model migration speed in response to changes in number of motors and clutches. Table 2 shows the parameters used in simulations. A unique prediction of the cell migration simulator is the optimality of the motor-to-clutch ratio (Fig 2E-F). If a cell is operating at the optimal motor-to-clutch ratio or has too few motors, decreasing motor activity will result in slower migration, as this directly decreases the cell’s ability to pull itself forward via detachment at the rear. However, if a cell has too many motors relative to the number of the clutches, a slight decrease in motor activity will allow the force to better distribute among the clutches, resulting in less clutch breakage, more efficient force transmission, and faster migration (Fig 2E). A similar effect can be seen in Figure 2F upon targeting of clutches. Of note, these simulations predict that cell migration would be more sensitive to decreasing motors, as a decrease in motors from the optimum has a very steep decrease from fast migration to no migration while a decrease in clutches from the optimum has a more gradual decrease in migration speed. This suggests that cell migration speed would be more sensitive to motor targeting drugs than to clutch targeting drugs.

**Table 2.**
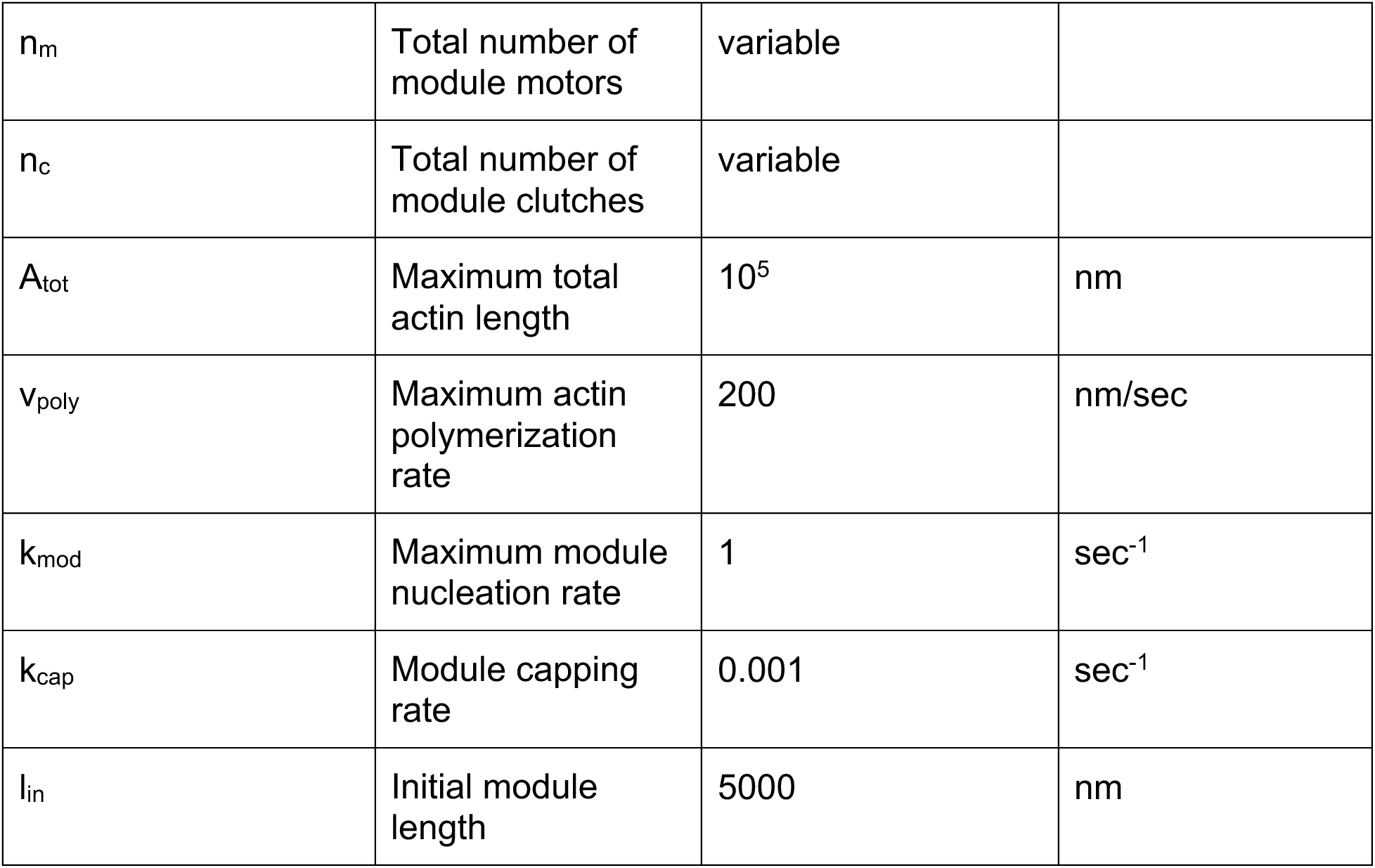
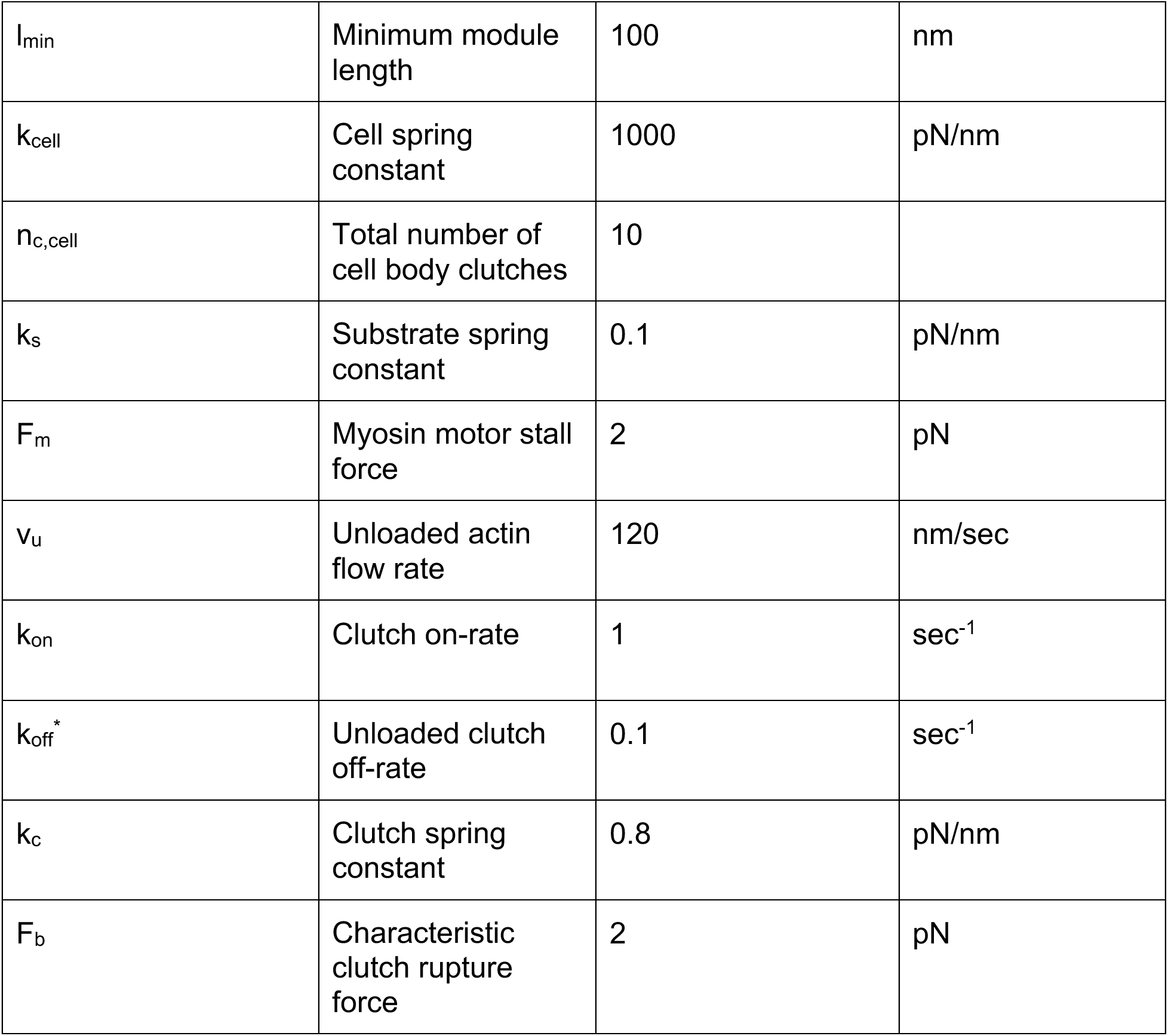
Cell Migration Simulator (CMS) parameters. All references values from Bangasser et al. [13].

|  |  |  |  |
| --- | --- | --- | --- |
| $n_m$ | Total number of module motors | variable | |
| $n_c$ | Total number of module clutches | variable | |
| $A_{tot}$ | Maximum total actin length | $10^5$ | nm |
| $v_{poly}$ | Maximum actin polymerization rate | 200 | nm/sec |
| $k_{mod}$ | Maximum module nucleation rate | 1 | $\text{sec}^{-1}$ |
| $k_{cap}$ | Module capping rate | 0.001 | $\text{sec}^{-1}$ |
| $l_{in}$ | Initial module length | 5000 | nm |
| $l_{\min}$ | Minimum module length | 100 | nm |
| $k_{\text{cell}}$ | Cell spring constant | 1000 | pN/nm |
| $n_{\text{c,cell}}$ | Total number of cell body clutches | 10 | |
| $k_s$ | Substrate spring constant | 0.1 | pN/nm |
| $F_m$ | Myosin motor stall force | 2 | pN |
| $v_u$ | Unloaded actin flow rate | 120 | nm/sec |
| $k_{\text{on}}$ | Clutch on-rate | 1 | $\text{sec}^{-1}$ |
| $k_{\text{off}}^*$ | Unloaded clutch off-rate | 0.1 | $\text{sec}^{-1}$ |
| $k_c$ | Clutch spring constant | 0.8 | pN/nm |
| $F_b$ | Characteristic clutch rupture force | 2 | pN |

### Glioma cell motility is biphasic with respect to myosin motor activity

After incubating green fluorescent glioblastoma cells (U251 and 6 PDX lines) with brain slices overnight, cells were imaged inside the slice for 10 hours. Figure 3A-B shows example tracks (wind rose plots) of single cell trajectories and Fig 3C-D shows the corresponding mean squared displacement (MSD) plots for this same subset of cells. The MSD plots, while variable, are generally linear and the cells therefore migrate via a random walk. Activity of myosin II motors can be decreased with blebbistatin treatment. Introducing various doses of blebbistatin into the brain slice assay leads to a progressive decrease in migration speed in U251 cells, with an over 4-fold decrease in random motility coefficient (RMC) at 250 μM blebbistatin (Fig 3E), implying that control U251 cells are sitting near or below the optimal motor-clutch level. At high doses of blebbistatin, all PDX lines exhibit a significant decrease in migration speed. At an intermediate dose, GBM85 + FBS, GBM 39, and GBM 117 exhibit an increase in migration speed (Fig 3F-G). This is a finding unique to a motor-clutch mode of migration, implying that these lines have too many motors relative to the number of clutches for optimal migration. By contrast, the osmotic engine does not utilize myosin, and so presumably operates in a manner independent of blebbistatin concentration. Each cell line has a unique estimated motor-to-clutch ratio predicted by response to motor targeting drugs and the ratio for each line relative to the optimum can be estimated based on these findings, as shown in Figure 3H. For example, GBM44 + FBS exhibits a progressive decrease in migration, implying that this line has either balanced motors and clutches or too few motors relative to the number of clutches, similar to U251 cells. By contrast, GBM80 + FBS, GBM85, and GBM12 exhibit no change in migration at intermediate doses, implying that they are slightly above the optimal motor to clutch ratio, so decreasing motors slightly pushes them over the optimum and we observe only modest change in migration speed, except at very high dose of blebbistatin. Lastly, GBM117, GBM39, and GBM85+FBS all exhibited a peak in migration at intermediate blebbistatin, consistent with their having a higher motor-to-clutch ratio under control conditions than the other cell lines. Supplementary Figure 1 includes the statistics for each curve. Altogether, we observed cell migration dynamics and sensitivity to motor perturbation consistent with a motor-clutch mode of migration for glioblastoma cells in brain tissue.

**Figure 3.**
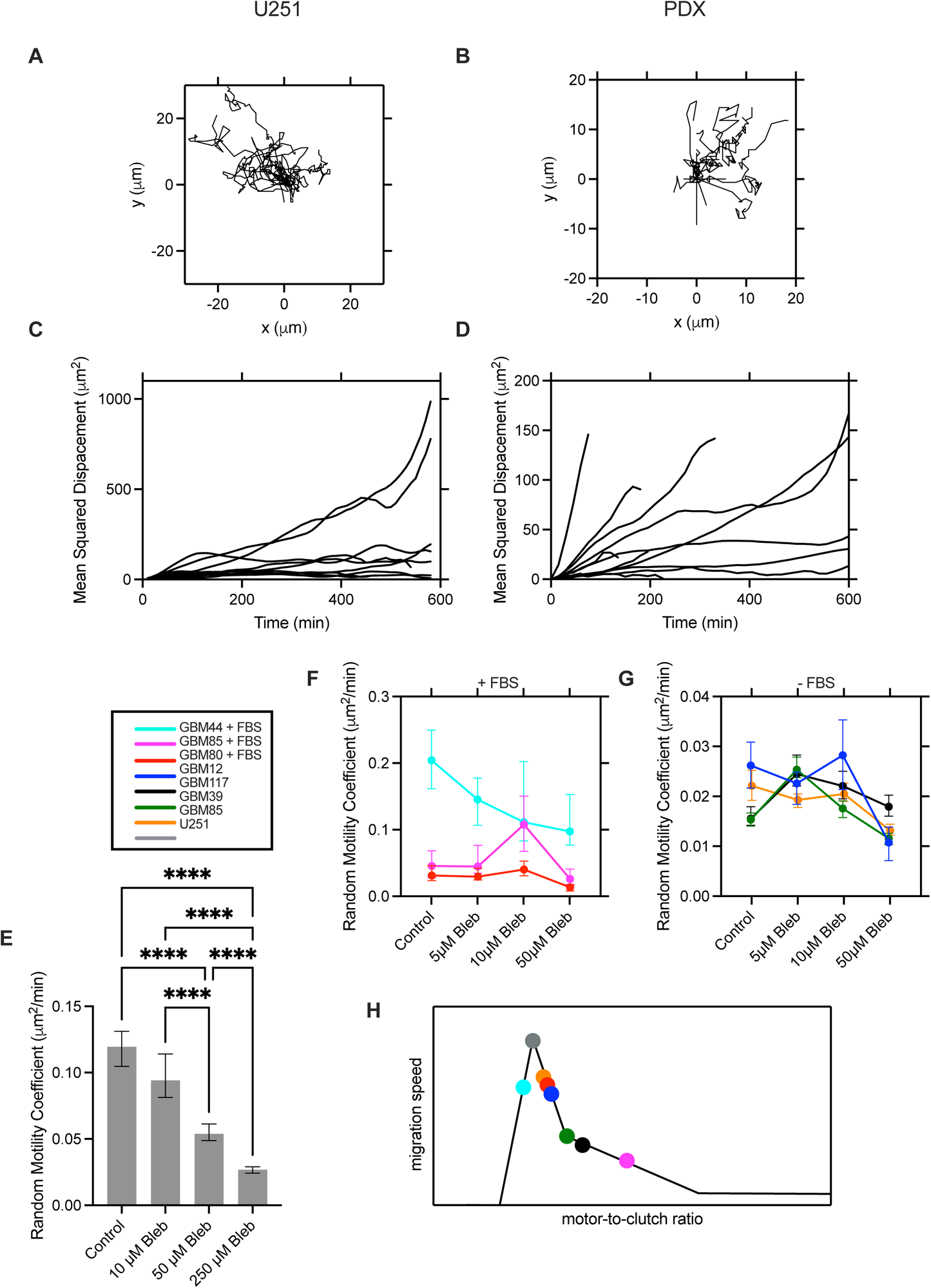
Migratory behavior of glioma cells in brain slices and response to myosin motor-targeting. Tracks of a) U251 cells and b) GBM80 (-FBS) PDX cells on brain slices. c-d) Corresponding mean squared displacement as a function of time interval for the same cells. e) RMC of U251 cells in mouse brain slices in response to increasing doses of blebbistatin. f-g) Response of PDX lines to varying doses of blebbistatin for cells that were cultured in the presence or absence of FBS. h) Simulated normalized migration speed vs. motor-to-clutch ratio with estimates motor-to-clutch ratio for each analyzed cell line.

### Integrins contribute to cell motility in brain tissue

Next we sought to determine the role of the clutch in cell migration. Initial studies examined migration on 2D polyacrylamide gels of various stiffnesses coated in various substrates to confirm the expected phenotype of a reduction in clutches. In this setup, the cell is forced to migrate using the specific clutch that binds to the ECM coating the gel. For example, on collagen I-coated PAG, cells migrate via integrins. In U251 cells, a knockout of talin1, an adaptor protein between integrins and F-actin, was developed using CRISPR/Cas9. These cells were unable to migrate on collagen gels (Fig 4A), indicating that integrin-mediated migration is nearly halted by knockout of talin1. In addition, the cells are smaller in projected area, as measured by a 65% decrease in projected area (Fig 4B), and more rounded, as measured by a 63% decrease in aspect ratio (Fig 4c), consistent with the hypothesis that the knockout of integrins prevents proper adhesion and cell spreading. U251 cells with talin1 knocked out migrated slower in brain tissue than controls by 3.5-fold (Fig 4D), though they were still migratory, indicating that integrin-mediated adhesion contributes to cell migration in brain tissue but is not the only mechanism the cells use to migrate.

**Figure 4.**
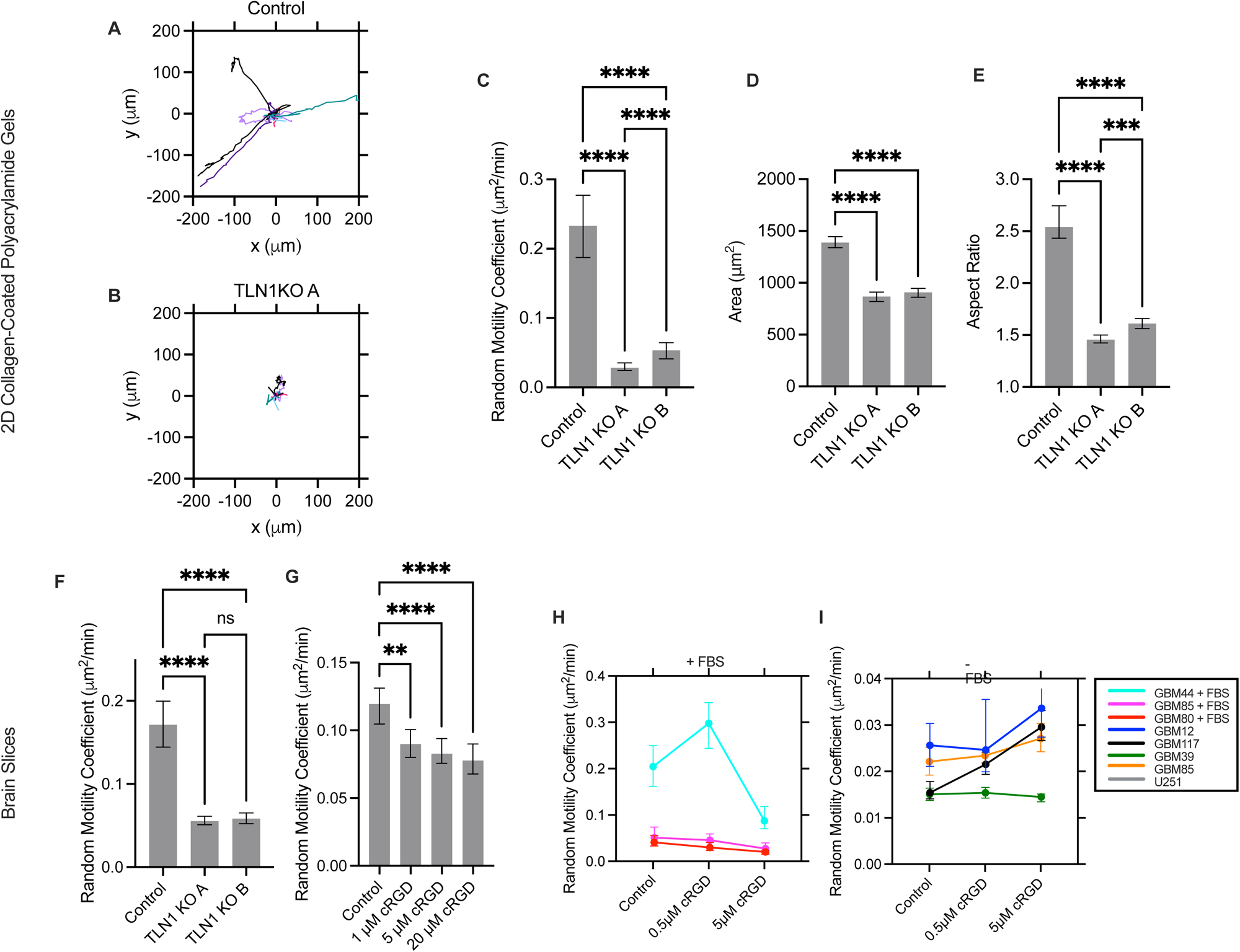
The role of integrins in GBM cell migration on PAGs and brain slices. Wind-rose plots for A) control cells and B) TLN1 KO cells. C) TLN1 KO cells migrate significantly slower than control lines on collagen-coated polyacrylamide gels. D) TLN1 KO cells are smaller in projected area and E) more round than control cells. F) TLN1 KO cells migrate slower on mouse brain slices. Inhibiting RGD-binding integrins with cyclo-RGD in G) U251 and H-I) PDX lines slows cell migration in some lines, while others are unaffected.

For both U251 and PDX lines, cyclo-RGD (cRGD) was used to target RGD-binding integrins (Fig 4E). This decreased migration speed in U251 by 1.5-fold, indicating that U251 cells are using RGD-binding integrins to migrate. In PDX lines (Figure 4F-G), some lines (GBM44+FBS, GBM85+FBS, GBM80+FBS, GBM39) exhibited a decrease in migration speed while GBM12, GBM85, and GBM117 did exhibit a significant effect. The statistics for this finding is summarize in Supplementary Figure 1. This indicates that while some lines do partially use integrins to migrate, others do not or can compensate for loss of RGD-binding integrins.

### CD44 contributes to cell motility

CRISPR/Cas9 was used to develop a knockout of CD44 in U251 cells. CD44 KO cells were extremely inefficient at adhering to α-CD44 mAb PAGs and the few cells that did adhere exerted lower traction force than control cells (Fig 5A). These cells also migrated slower in brain tissue with a 4-fold and 1.5-fold decrease in RMC observed in two knockout lines relative to control cells, indicating that both integrins and CD44 are important for cell migration in brain tissue (Fig 5B). When α-CD44 MAB was introduced into the brain slice assay with U251 cells, migration speed decreased by 1.5-fold, further indicating that CD44 is important for U251 migration (Fig 5C). PDX lines did not experience a decrease in migration speed in response to α-CD44 MAB incubation (Fig 5D). This is consistent with modeling results in Fig. 2 suggesting that cell migration is not particularly sensitive to clutch targeting. As these cells migrate slower than U251, a small decrease in migration speed would be difficult to detect.

**Figure 5.**
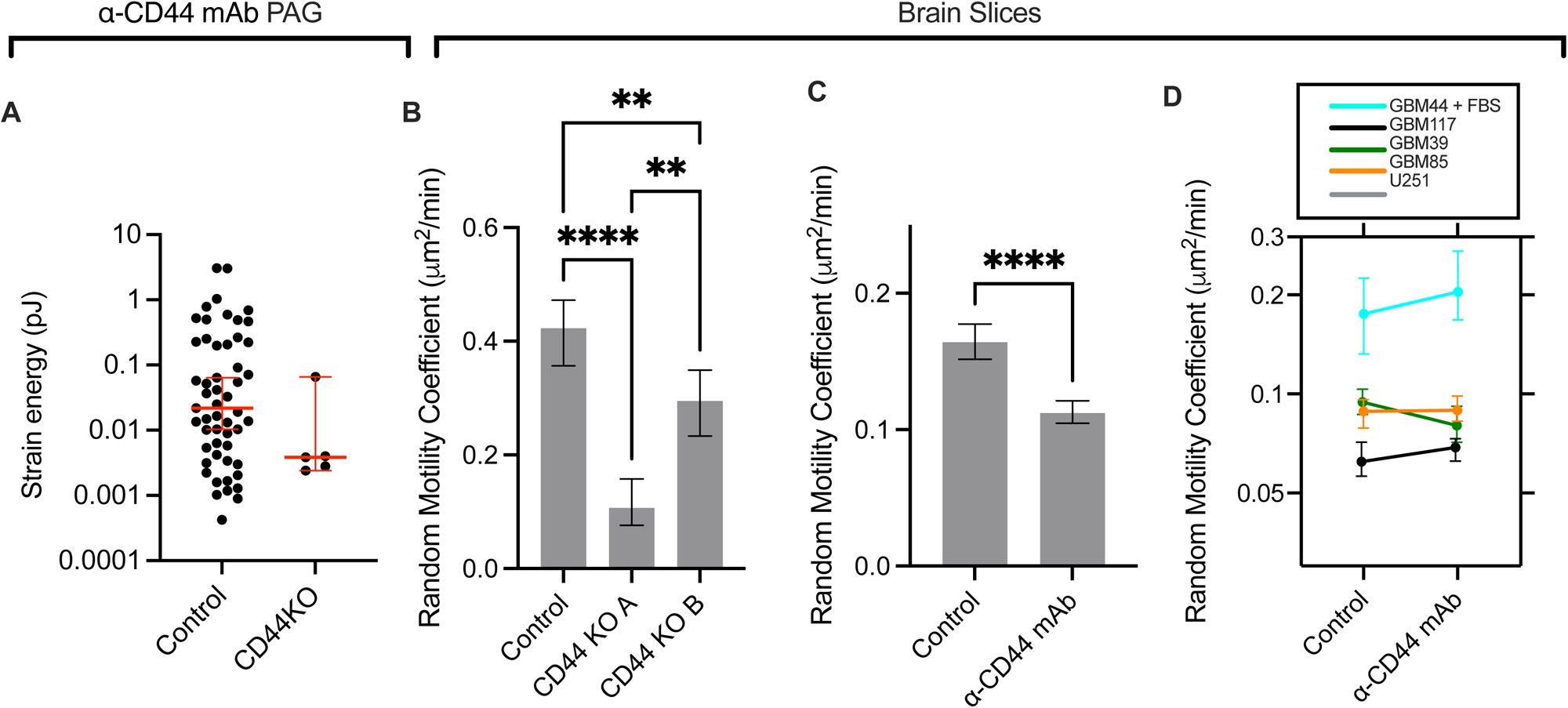
The role of CD44 in GBM cell migration on brain slices. A) CD44KO cells experience decreased adhesion to and lower traction strain energy on polyacrylamide gels coated in α-CD44 mAb. B) CD44 knockout cells migrate slower on brain slices relative to the control line. C) U251 cells on brain slices migrate slower with α-CD44 mAb incubation. D) Migration of PDX lines is unaffected by α-CD44 mAb incubation.

### Additive effects of simultaneous targeting of CD44 and integrins

In order to determine the interplay between CD44 and integrins, we targeted both integrins and CD44 simultaneously. When cRGD was used on the talin1 KO cells it had no impact on migration speed in brain tissue (Figure 6A), indicating that targeting integrins via talin knockout and RGD targeting drugs is not additive. On the other hand, cRGD was used on the CD44 KO cells and this further decreased migration in brain tissue (Figure 6B). When combined, there was a 10-fold decrease in RMC in cRGD targeting of CD44 KO cells relative to control, suggesting that targeting of CD44 and integrins is additive. In wild-type U251 cells, simultaneous targeting of α-CD44 mAb and cRGD further decreased migration speed compared to either condition alone with a 2.3-fold decrease relative to control (Figure 6C). These results are consistent with the finding that glioma cells utilize both integrins and CD44 as clutches independently to migrate. While targeting one may be able to partially decrease migration, both integrins and CD44 must to be targeted to observe a large decrease in migration speed.

**Figure 6.**
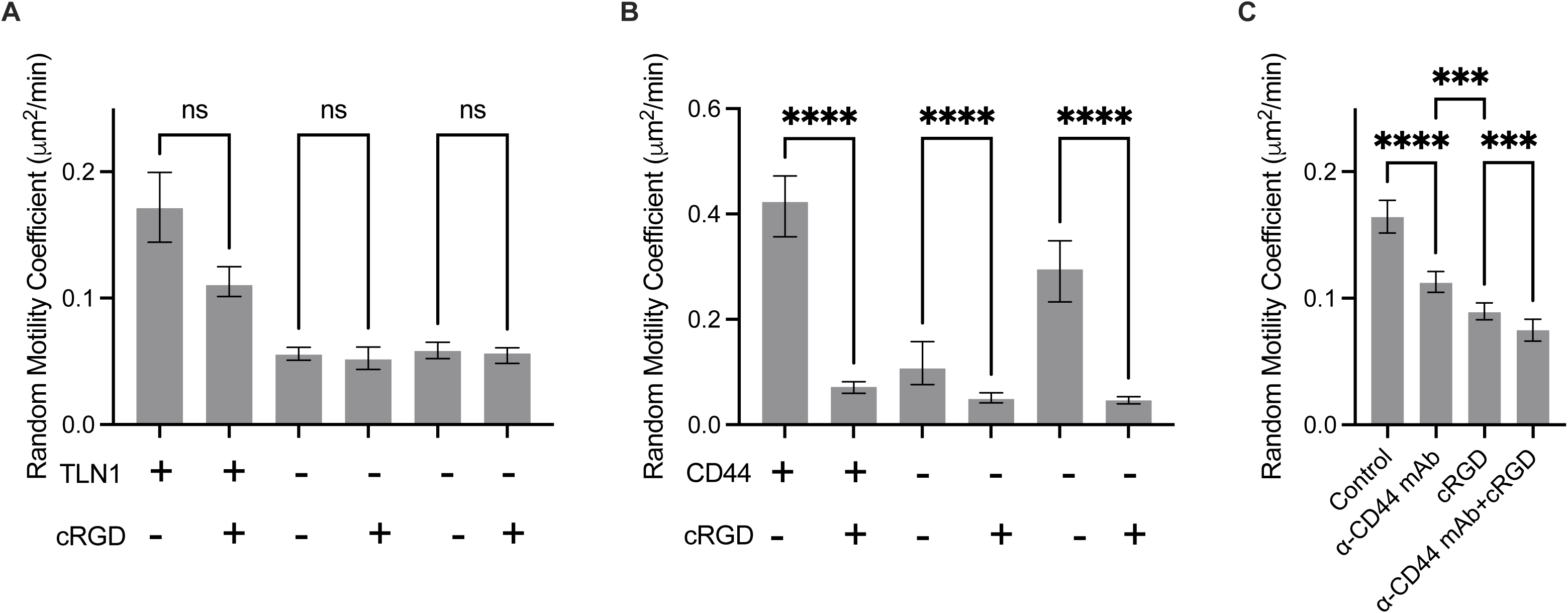
Combined targeting of integrins and CD44 maximally inhibits migration on brain slices. A) cRGD targeting of TLN1 KO cells does not further decrease migration. B) cRGD targeting of CD44KO cells decreases migration further on brain slices. C) Combination targeting of U251 cells on brain slices with α-CD44 mAb and cRGD decreases migration more than either by itself.

## Discussion

In this study, we utilized imaging of GBM cell migration to identify the mechanism of single-cell migration in brain tissue for both high passage U251 cells and low passage patient-derived xenograft lines. Confocal imaging of green-fluorescent cancer cells on red-fluorescent vasculature revealed the cells pulling on the vasculature at the leading edge, a prediction unique to a motor clutch mode of migration and contradictory to an osmotic engine mode of migration. In addition, blebbistatin was used to target myosin II motors and demonstrated the significance of motors to cell migration. Further, some PDX lines were found to have a biphasic response to blebbistatin as described by an increase in migration speed at intermediate doses, another finding that is a specific prediction of a motor clutch model. Through knockouts and drug targeting, both integrins and CD44 were found to act as clutches and the slowest migration was achieved by targeting both clutches simultaneous. This is significant because it both describes a mechanism for how these cells are migrating and suggests that the optimal way to treat GBM via targeting cell migration would be to target both integrins and CD44 simultaneously.

There are a few notable limitations to the methods used. In analyzing the environmental mechanical deformations, many cells showed no visible interaction with the environment, especially in the PDX lines, which could be due to a number of reasons. For example, the cells could be adhered to ECMs that are not visualized by the microscope such as ECMs associated with white matter tracts. They could also simply be non-migratory or the deformation is too small for the microscope to resolve, which is likely the case in the PDX lines as those are slower migrating. In addition, televascular cell-ECM interactions were not imaged and could behave differently than cell-vasculature interactions.

Despite these limitations, the present findings are significant because the prediction of cells pulling on vasculature is unique to a motor clutch mode of migration, and of the cells whose interaction with vasculature could be visualized all but one demonstrated pulling at the leading edge. This is very strong evidence that these cells use a motor clutch mode of migration while also arguing strongly against the competing osmotic engine model. Had the cells been migrating using an osmotic engine, pushing at the leading edge would have been observed. The hypothesis that cells migrate using a motor-clutch mode of migration was further supported by targeting of motor-clutch components. Most significantly, some lines were found to have a biphasic response to blebbistatin treatment. This optimality of motors and clutches is a also unique prediction of the motor-clutch model. While it’s possible that blebbistatin treatment could indirectly impact migration under an osmotic engine model, there is no obvious mechanism for a biphasic response to blebbistatin in this model. Further, as adhesions purely act as frictional resistance to migration in an osmotic engine model, the decrease in migration in response to targeting multiple clutches through a variety of mechanisms argues against the osmotic engine model and supports the motor-clutch model.

While a previous study examined a limited number of U251-mediated vasculature deformations in brain tissue *ex vivo* [22], traction analysis in brain slices of low passage glioblastoma lines had not been done previously. The low-passage PDX cells behaved similarly to U251 cells, though they generally migrated slower overall. The present study analyzed PDX lines with and without media containing fetal bovine serum per Mayo PDX culture protocol, and noted significant differences between the two conditions. It was consistently found that lines cultured in serum exhibited significantly faster migration, including GBM85 which was studied in both serum and serum-free conditions. FBS therefore has some activating effect on cell migration in the PDX lines. Serum exposure is not an unrealistic experimental effect, as tumors experience breakdown of the blood-brain barrier that presumably exposes tumor cells to serum. Thus, we speculate that the breakdown of the blood-brain barrier in the central regions of the tumor [57] could provide an impetus for accelerating glioblastoma cell migration, which presumably would enhance cell invasion into adjacent relatively normal brain parenchyma where the vasculature barrier function is still intact.

Our study also found that for U251 cells, the slowest migration was achieved by dual targeting of integrins and CD44, suggesting that both molecules are important clutches. This implies that in developing clutch-targeting treatments, targeting both mechanisms of migration simultaneously will have the greatest effect. In the PDX lines, α-CD44 mAb incubation did not decrease migration speed as hypothesized. This could be because the cells migrate so slowly that detecting a partial decrease in migration speed is not feasible, as results from the CMS simulations indicate that migration is less sensitive to targeting of clutches than motors. This further suggests that targeting one clutch at a time may not be sufficient to adequately slow down migrating cells. It is also possible that the PDX lines do not use CD44 as a clutch, however previous findings strongly suggest a role of CD44 in glioblastoma migration [37]. An alternative strategy would be to target myosin II motors (Picariello et al., PNAS, 2019), which the CMS predicts is very sensitive (Fig. 2e).

In summary, our study investigated a variety of glioblastoma cell lines ranging from low to high passage in an environment that closely mimics cells in the invasive front of a brain tumor. We find that treatment of glioblastoma via targeting migration should either simultaneously target both integrins and CD44, or else target myosin II. This study also highlights the importance of determining whether cells are at an optimal motor-clutch ratio, as insufficient targeting of motors or clutches can potentially increase migration. The motor-clutch-based CMS predicts that the former is more sensitive, and may be a better strategy, although it will be important to assess the effects of drugs on immune cells in the glioblastoma tumor microenvironment, especially antitumoral CD8+ T cells. If the drugs impair T cells to a similar or greater extent than they impair glioblastoma cells, then there will be no benefit or potentially harm to the patient. Overall, our study provides a framework for further development of antimigratory strategies in glioblastoma, and potentially other invasive cancers as well.

## Acknowledgements

Research reported in this publication was supported by NIH grants U54CA210190, P01CA254849, and U54CA268069. We thank members of the Odde lab for helpful conversations throughout the course of this work. The content of this work is solely the responsibility of the authors and does not necessarily represent the official views of the NIH.

## Author Contributions

MK performed the traction force experiments on U251 CD44 KO cells. SMA completed the rest of the work in the paper.

## Supplementary Figures

**Supplementary Figure 1.**
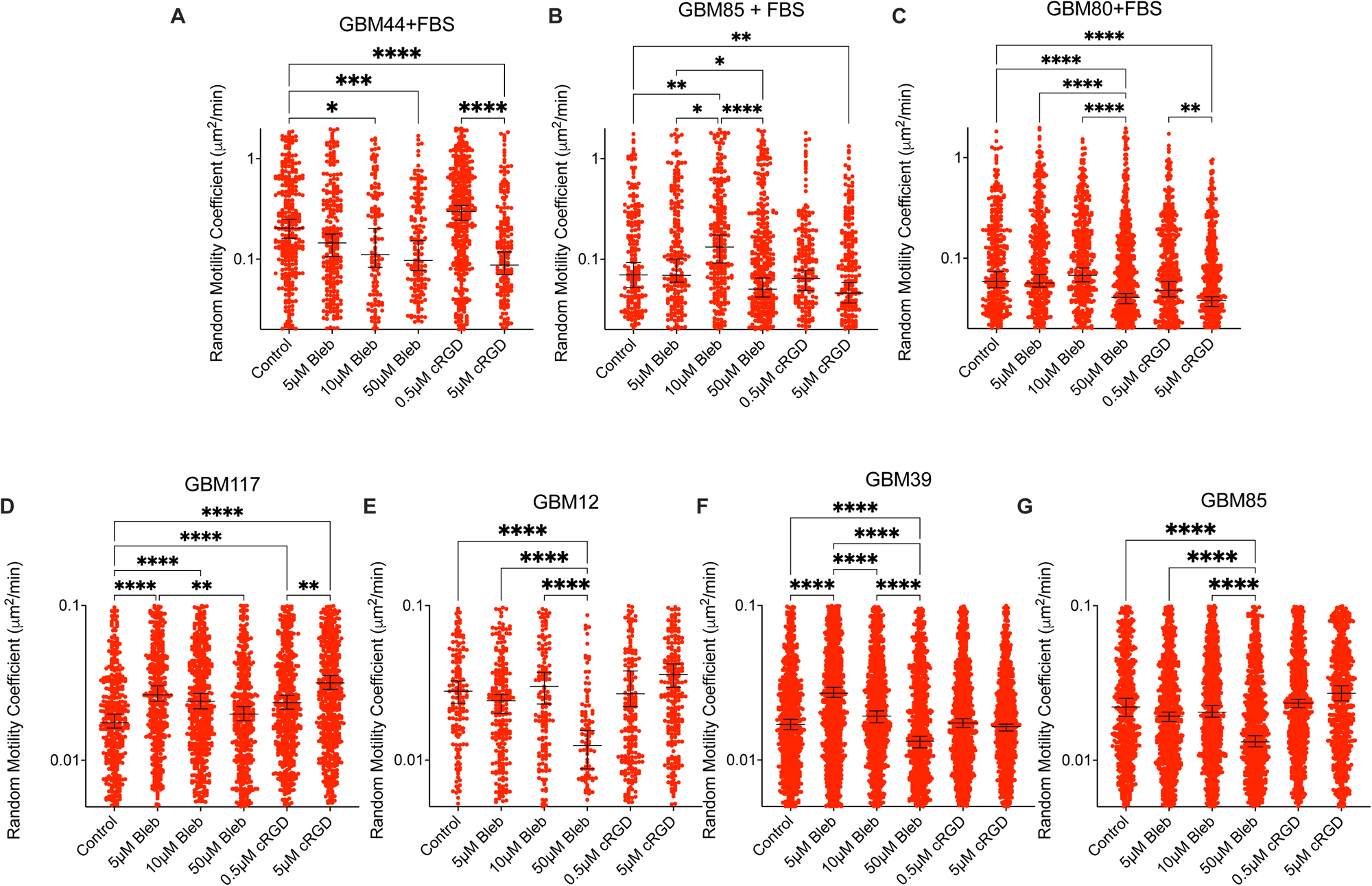
Migratory behavior of individual glioma cells in brain slices in response to blebbistatin and cyclo-RGD. RMC of cells in response to varying doses of blebbistatin and cRGD for A) GBM44 + FBS, B) GBM 85 + FBS, C) GBM 80 + FBS, D) GBM 117, E) GBM 12, F) GBM 39, and G) GBM 85. Each dot represents a single cell. The black line denotes the median value and 95% confidence interval. Statistics were done using a Kruskal Wallis test.

**Supplementary Figure 2.**
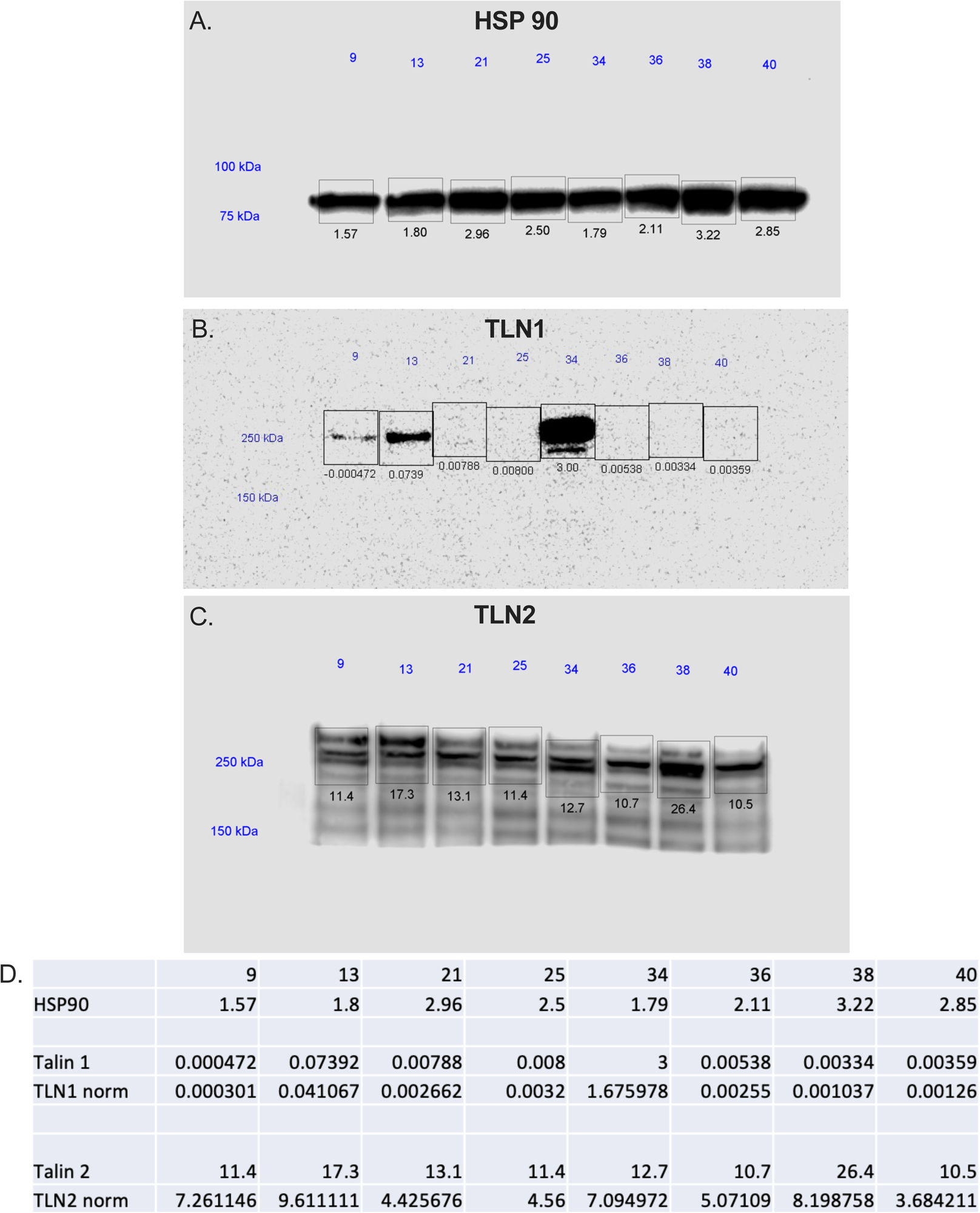
Western Blot demonstrating knockout of talin 1. Western Blot results for the A) control HSP90, B) Talin1, and C) Talin2 in development of Talin1 knockout. D) Calculations demonstrating knockout of talin1 in given samples and no significant upregulation of talin 2.

## Supplementary Movie Captions

Supplementary Movie 1. PDX cell pulls on vasculature. Example movie of GBM44 (green) pulling on vasculature (magenta) as it migrates.

Supplementary Movie 2. PDX cell pulls on vasculature. Example movie of GBM44 (green) pulling on vasculature (magenta) as it migrates.

Supplementary Movie 3. U251 cell pulls on vasculature. Example movie of U251 (green) pulling on vasculature (magenta) as it migrates.

Supplementary Movie 4. U251 cell pulls on vasculature Example movie of U251 (green) pulling on vasculature (magenta) as it migrates.

Supplementary Movie 5. U251 cells migrating in brain tissue. Example movie of U251 cells (green) migrating in brain tissue with vasculature and microglia (magenta).

Supplementary Movie 6. PDX cells migrating in brain tissue. Example movie of GBM80 cells (green) migrating in brain tissue with vasculature and microglia (magenta).

## Notes

### Competing Interest Statement

The authors have declared no competing interest.

